# Local translation of circular RNAs is required for synaptic activity and memory

**DOI:** 10.1101/2025.09.18.676988

**Authors:** Hao Gong, Haobin Ren, Wei-Siang Liau, Qiongyi Zhao, Alexander P. Walsh, Mason R.B. Musgrove, Joshua W.A. Davies, Esmi L. Zajackowski, Sachithrani U. Madugalle, Paul R. Marshall, Ziyue Xu, Tian Lan, Jichun Shi, Jiazhi Jiang, Wei Wei, Xiang Li, Irina Voineagu, Juli Wang, Sania Sultana, Chun-Kan Chen, Lingrui Zhang, Sourav Banerjee, Victor Anggono, Howard Y. Chang, Timothy W. Bredy

## Abstract

Circular RNAs (circRNAs) are enriched in synapses and implicated in cognitive processes, and recent studies have shown that circRNAs can encode micropeptides, which suggests that there may be novel synaptic proteoforms in the brain that await discovery. Here we report widespread learning-induced local circRNA translation in the prefrontal cortex of male C57BL/6 mice. More than 1500 synapse-enriched circRNAs contain active IRES elements, with 842 interacting with the ribosome and 241 exhibiting direct evidence of activity-induced translation. We discovered a synapse-enriched micropeptide (P1) that is derived from a single exon circRNA, the mRNA host of which encodes an enzyme associated with protein repair. Although P1 is only a third of the size of the full-length protein, it is locally expressed, enzymatically active, and interacts with plasticity-related proteins, including CaMKIIα. In addition to direct effects on synaptic activity, targeted P1 knockdown impairs whereas its overexpression enhances fear extinction memory. These findings shed new light on the ‘dark’ proteome in the brain and reveal local circRNA translation as a novel mechanism of plasticity and memory.

## Introduction

Circular RNAs (circRNA) belong to a structurally distinct family of regulatory RNAs comprised of single-stranded closed-loop RNA molecules, which are highly resistant to degradation^1^. They are abundant in the brain and enriched within synapses^2,3^, with emerging studies suggesting a relationship between circRNA dysregulation and cognitive impairment^4–6^. In line with these observations, we recently discovered a role for synapse-enriched circRNAs in fear extinction memory^7^.

It has become evident that many circRNAs can undergo cap-independent translation to generate functional micropeptides that are involved in a variety of cellular processes^8,9^. These findings suggest that there may be a significant population of novel proteoforms in the brain that have yet to be revealed, which led us to question whether there are synapse enriched circRNA-derived peptides that may serve to regulate memory. Fear extinction is an important form of associative learning that is supported by coordinated changes in gene expression, protein synthesis, and the activity of regulatory RNAs within the medial prefrontal cortex (mPFC)^10–13^. Although some of the basic mechanisms of this cognitive process have been revealed, the complete molecular code underlying the formation of fear extinction memory remains to be fully determined.

To address this question, we used a combination of high-throughput Internal Ribosome Entry Site (IRES) reporter assays, ribosome sequencing, and mass spectrometry on synaptosomes isolated from the mPFC of fear extinction-trained mice to examine learning-induced circRNAs in the synapse. We discovered many have features of translational capacity, including one such circRNA-derived micropeptide, we call P1. This novel synapse-enriched micropeptide is catalytically active, interacts with a variety of plasticity-related synaptic proteins, including CaMKIIα, influences the frequency of spontaneous mEPSCs in cortical neurons, and is required for fear extinction memory. CircRNAs have therefore evolved to overcome the challenge of space and time in neurons, in part, by establishing a localised network of circRNA-derived micropeptides that function in a state-dependent manner to control synaptic plasticity and memory.

## Results

### Identification of translationally active circRNAs at the synapse

To begin to explore the possibility of learning-induced translation of circRNAs in the synaptic compartment, we examined our previously published synapse-enriched circRNA-seq data^7^, which revealed 1,819 circRNAs associated with both fear-conditioning (retention control, RC) and fear extinction learning (EXT) in the mPFC in adult male C57Bl/6J mice (Fig. 1a and Supplementary File 1). 91.6% of synapse-enriched circRNAs were derived from exons, which is suggestive of their potential to be translated (Supplementary File 1). Full-length isoform prediction analysis indicated that the majority of synapse-enriched circRNAs comprise 2 or 3 exons (∼37% and ∼32%, respectively), with most synapse-enriched circRNAs spanning 100 to 450 nucleotides in length (∼80.6%) (Fig. 1b and 1c, Supplementary File 2), which perhaps indicates a high level of stability of circRNAs with shorter length and fewer exons in the synaptic compartment.

**Figure 1.**
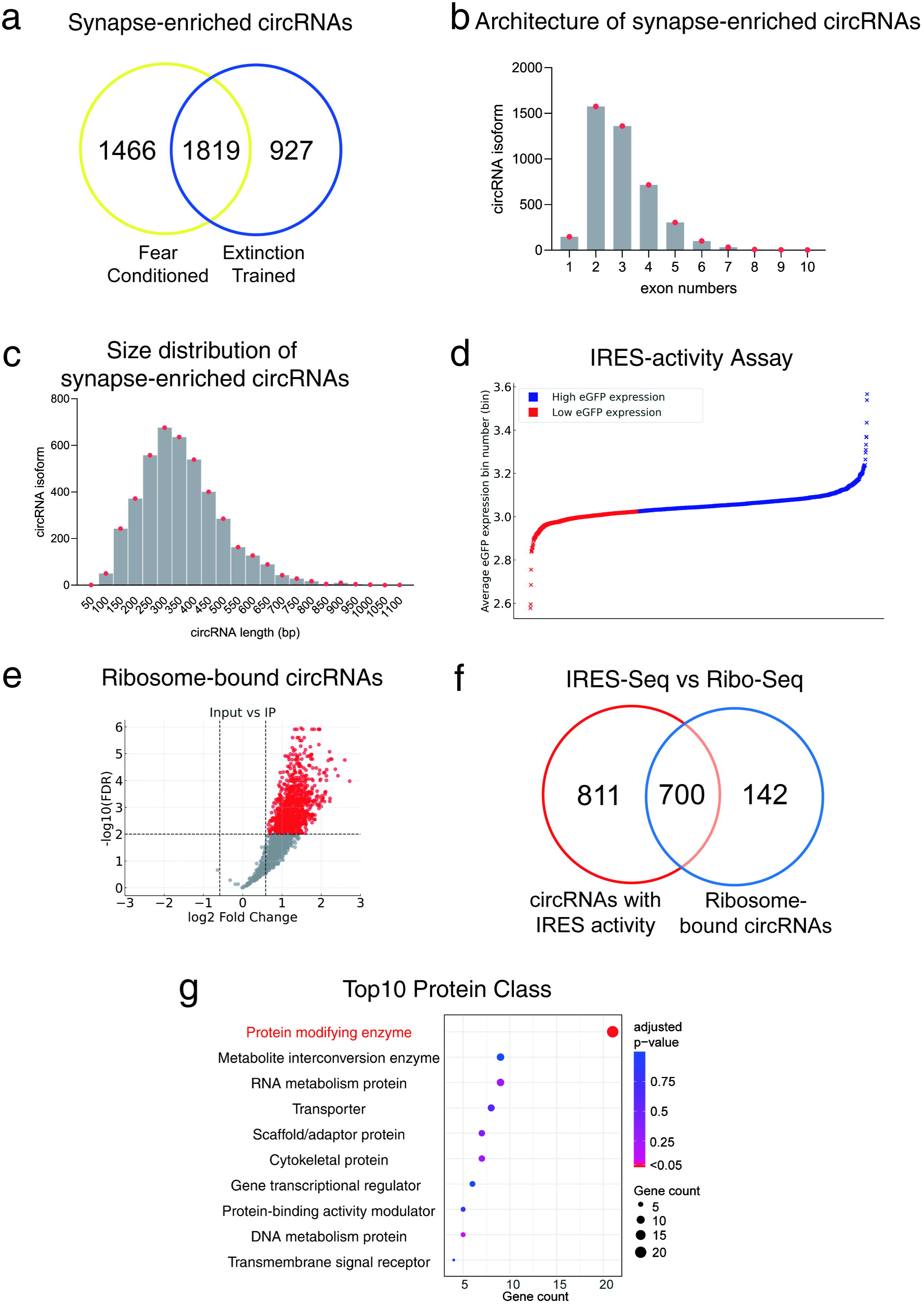
**a.** Venn diagram representing 1,819 synapse-enriched circRNAs that are significantly increased in both fear conditioning and extinction trained mice. **b.** Distribution of circRNA isoform exon numbers, **c.** Size distribution of circRNA isoforms. Both plots are derived from EXT-trained animal mice with full-length circRNA prediction applied to the 1,819 synapse-enriched circRNAs in panel a, of which 1,098 circRNAs (4,250 isoforms) were available. **d**. eGFP expression reflecting IRES activity. A total of 1,511 out of 1,819 captured circRNA isoforms were identified with eGFP expression scores divided by K-means clustering. The high eGFP expression group had 1033 (eGFP expression [bin] = 3.02-3.56) and the low eGFP expression group had 478 (eGFP expression [bin] =2.57-3.02). **e.** Volcano Plots for 1,450 synapse-enriched circRNAs bound to the ribosome. This volcano plot illustrates Input vs Rpl22-HA-IP. Each dot represents an individual circRNA isoform, with red dots indicating ribosome-bound circRNAs (FDR < 0.05, Log₂FC > 0.585, n=842), and gray dots represent all other detected circRNAs (n=608). **f.** Venn diagram representing the number of circRNAs that exhibit both IRES activity and are bound to the ribosome. **g.** GO analysis of the top 10 protein classes of parent genes from which circRNAs were found to generate unique peptides. Protein classes with an adjusted p-value < 0.05 are shown in red, while classes with an adjusted p-value ≥ 0.05 are represented in shades of blue.

We next employed a high-throughput IRES-reporter screen to identify synapse-enriched circRNAs exhibiting cap-independent translation^9^. We designed 3,139 synthetic oligonucleotides spanning the back splice junction (BSJ) of the differentially expressed circRNAs described above and cloned them into an oligo-split-eGFP circRNA reporter construct (Supplementary Fig. 1a, 1b and Supplementary file 3). The construct pool was then transfected into HEK293T cells and cultured for 5 days, *in vitro*. Fluorescence-activated cell sorting (FACS) was used to isolate cells expressing eGFP followed by high-throughput DNA sequencing to quantify translation activity. The mean weighted rank distribution of reads across the bins was used to quantify eGFP expression for each putative IRES element, revealing 1,511 circRNAs that exhibit translation potential (Fig. 1d, Supplementary File 4).

Encouraged by the observation of abundant IRES activity of synapse-enriched circRNAs, we next asked whether there was any evidence of *in vivo* experience-dependent circRNA translation using a transgenic mouse called Tagger that enables neuron-specific enrichment of translating RNA, which we previously used to examine mechanisms of memory^14,15^. Naïve 9-week-old male CaMKIIα Cre-Tagger mice (n=12/group) were implanted with a 22-gauge dual guide cannula into the mPFC, after which they were given 1 week to recover. The mice underwent fear training with 3 pairings of a tone (2 min, 80 Hz) with a foot shock (0.7 mA, 2 s). On Day 8, they received extinction training with 60 non-reinforced tone exposures (EXT) and were sacrificed immediately after fear extinction training and mPFC tissue prepared for analysis. Following the isolation of synaptosomes by fractionation, RNA immunoprecipitation for potentially translating RNAs was performed using Rpl22-HA pull down, and 200ng of barcoded RNA was prepared for sequencing (Ribo-Seq). The synapse-enriched circRNAs identified above were then overlayed with the Ribo-seq data (Supplementary File 5), revealing 842 synapse-enriched circRNAs that show increased interaction with the ribosome in response to learning (Fig. 1e), 700 of which also overlapped with the IRES activity assay (Fig 1f). These findings suggest that a substantial number of synapse-enriched circRNAs are either involved in the translation of other mRNAs or can generate their own peptides.

To differentiate between these two possibilities, we examined the predicted full-length sequence of the ribosome bound circRNAs and identified 61 that contained a “GGUGGU” motif, which is an RNA sequence that belongs to the B2 family of SINE retrotransposons (Supplementary File 6). Recent work has shown that noncoding RNAs that comprise an inverted SINEB2 sequence can promote the translation of target mRNAs^16^. Importantly, circRNAs have also recently been identified with this “SINEUP” capacity^17^. It is possible that when a circRNA interacts with the ribosome, it need not generate peptide but, instead, may have evolved the capacity to promote the translation of mRNAs that are coincidentally bound at the same time. Furthermore, we also found that 213 ribosome-bound circRNAs contain a “GUGGC” motif (Supplementary File 6), which is the smallest catalytic RNA sequence identified to date^18^. Although speculative at this time, given that ribozyme activity has recently been discovered in the brain and shown to be involved in learning and memory^19^, this observation raises the interesting possibility that synapse-enriched circRNAs harbouring this catalytic sequence motif may potentially function as ribozymes.

To provide direct support for the idea that there is learning-induced circRNA translation at the synapse, we examined synaptosome peptide fragments in samples derived from fear extinction-trained mice using High-Performance Liquid Chromatography-Tandem Mass Spectrometry (HPLC-MS/MS). Open reading frame (ORF) prediction (TransDecoder, v5.5.0) was used to identify putative small ORFs derived from synapse-enriched circRNAs, which were then cross-referenced with the MS data. We found 1,434 circRNAs containing MS-matched tryptic peptides that overlapped with predicted ORFs with 95% confidence, and 241 unique MS-matched tryptic peptides spanning the circRNA BSJ (Supplementary File 7). These findings provide strong evidence for learning-induced generation of novel circRNA-derived peptides in the synaptic compartment. Next, a gene ontology (GO) analysis on the parent circRNA protein-coding host genes revealed significant enrichment for proteins related to enzymatic activity, which included 21 circRNA-derived peptides (Fig. 1g). Based on these findings, we selected several for further consideration, including *circHecw1* and *circMycbp2* (whose host proteins exhibit E3 ubiquitin ligase activity), *circPak1* (whose host is a serine-threonine protein kinase involved in Rac1 signaling), *circUxs1* (whose host is a relatively unknown UDP-glucuronate decarboxylase), and the functionally uncharacterized circRNA, *circPcmtd1*. This circRNA attracted our attention because it is derived from a single exon and its host gene *Pcmtd1* (protein-L-isoaspartate O-methyltransferase domain containing one) is a linear multi-exon mRNA that is a homologue of the protein repair enzyme, Protein L-isoaspartyl/D-aspartyl O-methyltransferase (PIMT). Importantly, PIMT is critically involved in learning and memory^20^ and has been shown to be neuroprotective in models of age-related neurodegenerative disorders^21^.

### *CircPcmtd1* generates a novel synaptic micropeptide

*CircPcmtd1* is generated through back-splicing of a single exon from the multi-exon Pcmtd1 transcript (Supplementary Fig. 2a and b) and is highly expressed in response to learning (Supplementary Fig. 2c and d). Having demonstrated that synapse-enriched *circPcmtd1* interacts with the ribosome, contains an IRES with strong translation activity and generates a novel micropeptide (which we have named “P1”), we next sought to better understand its protein architecture. Relative to linear Pcmtd1 mRNA, the start codon in exon 2 did not change following circPcmtd1 back-splicing; however, a new stop codon was generated, which resulted in a unique peptide (Supplementary Fig. 2a and Fig. 2a). Pfam analysis of the predicted P1 sequence indicated that it is in fact a truncated form of Pcmtd1 and belongs to the PIMT family (Fig. 2a). An examination of the P1 amino acid sequence revealed a simplified protein comprised of 137 amino acids, which is well within the range of being denoted as a micropeptide. P1 contains a highly conserved S-adenosyl-L-methionine-dependent methyltransferase domain and a unique intrinsically disordered region (IDR) spanning the BSJ of circPcmtd1 (Fig. 2a and 2b). The *de novo* generation of a novel IDR indicates potential differences in folding, kinetics, and protein-protein binding affinities.

**Figure 2.**
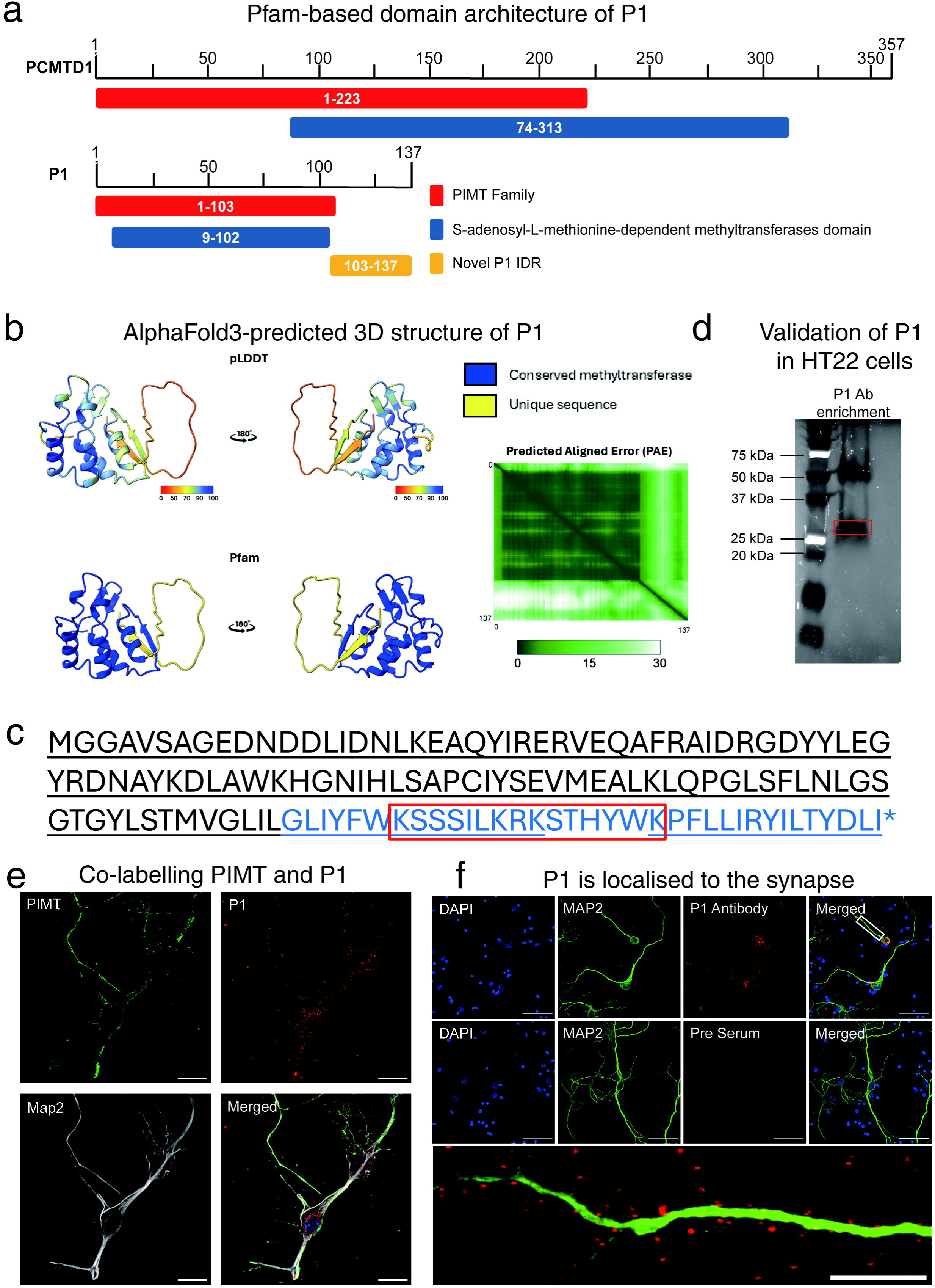
**a.** A schematic of Pfam analysis of the protein coding-host locus Pcmtd1 and P1. **b.** AlphaFold3 structure prediction of P1. The left panel shows the Pfam domain annotation, where the blue region indicates a conserved methyltransferase domain, and the yellow region represents a unique IDR sequence specific to P1. The right panel displays the pLDDT (predicted Local Distance Difference Test) confidence scores across the predicted amino acid sequence, color-coded from red to blue. Blue corresponds to high-confidence structural regions, while red indicates low-confidence regions, which are often associated with IDRs. Predicted Aligned Error (PAE) reflects the model’s confidence in the relative positioning of residue pairs. **c.** The custom-made antibody directed against P1 is highlighted by in red. The IP-MS detected fragments using this antibody in neurons is highlighted by underline. The unique peptide of P1 is highlighted in blue font. **d.** SDS-PAGE showed the size of P1 detected in HT22 cells. Red box, the extracted region for IP-HPLC-MS/MS. Only the 26kDa region was revealed the P1 sequence. **e.** Immunofluorescence detection shows P1 has unique distribution pattern relative to its homologue PIMT in primary cortical neurons. Confocal images show the expression of P1 and PIMT along dendrites in primary cortical neurons (MAP2). For left to right and top to bottom, the images show P1 (green), PIMT (red), and the MAP2 (white). Scale bars, 50µm. **f.** Confocal images of neurons show synaptic expression of P1 along MAP2 labelled dendrites in primary cortical neurons. Images on the left show DAPI labelling; the middle panels label the morphology of neurons (anti-MAP2 primary antibody), and red dye labels the P1 protein (the top image used customized P1 antibody, and the bottom image is negative control). The bottom panel shows an enlarged region of dendrites. Scale bars, 50µm, 10µm (enlarged image).

To provide further evidence for endogenous P1 micropeptide expression, we generated a custom antibody directed against the unique IDR and performed immunoprecipitation HPLC-MS/MS on primary cortical neurons (IP-), *in vitro*, which revealed tryptic peptides aligned with the predicted amino acid sequence, including the unique IDR region of P1 (Fig. 2c). In addition, a strong signal corresponding to the correct predicted size of P1 (∼26 kDa) was observed by western blot in HT22 cells (Fig. 2d). Co-labeling in primary cortical neurons, *in vitro*, revealed that PIMT is primarily membrane associated whereas P1 is cytoplasmic and enriched at the synapse, which suggests that there may be specific targets for P1 that are distinct from those of PIMT (Fig. 2e). Furthermore, we also observed robust punctate P1 protein labeling along dendrites, strongly indicative of synaptic localization (Fig. 2f).

### P1 interacts with synaptic proteins that are required for memory formation

To elucidate the potential underlying mechanism of how P1 might regulate localized synaptic plasticity and memory, we performed immunoprecipitation using the custom P1 antibody on samples derived from the mPFC of behaviorally trained mice, followed by HPLC-MS/MS. 77 proteins, including a number of neuron-specific proteins involved in synaptic plasticity and memory such as growth associated protein 43 (Gap-43, also known as neuromodulin)^22^; neural cell adhesion molecule 1 (Ncam1), a synaptic protein shown to be involved in memory across several species^23^; limbic system-associated protein (Lsamp), which is another neural adhesion molecule that is critically involved in synaptogenesis^24^; autism-related synaptotagmin-like protein 4 (Sytl4)^25^, and the key plasticity-related protein calcium-calmodulin (CaM)-dependent protein kinase II alpha (CaMKIIα)^26^ were found to directly interact with P1 (Fig. 3a and Supplementary File 8). In fact, amongst the proteins that were observed to interact directly with P1, CaMKIIα was exclusively detected in the extinction learning group (Fig. 3a). CaMKIIα was intriguing because it is activity-dependent, known to aggregate in the synaptic compartment, perpetuates its own activity state at the synapse, and is directly involved in plasticity related to learning and memory^27,28^.

**Figure 3.**
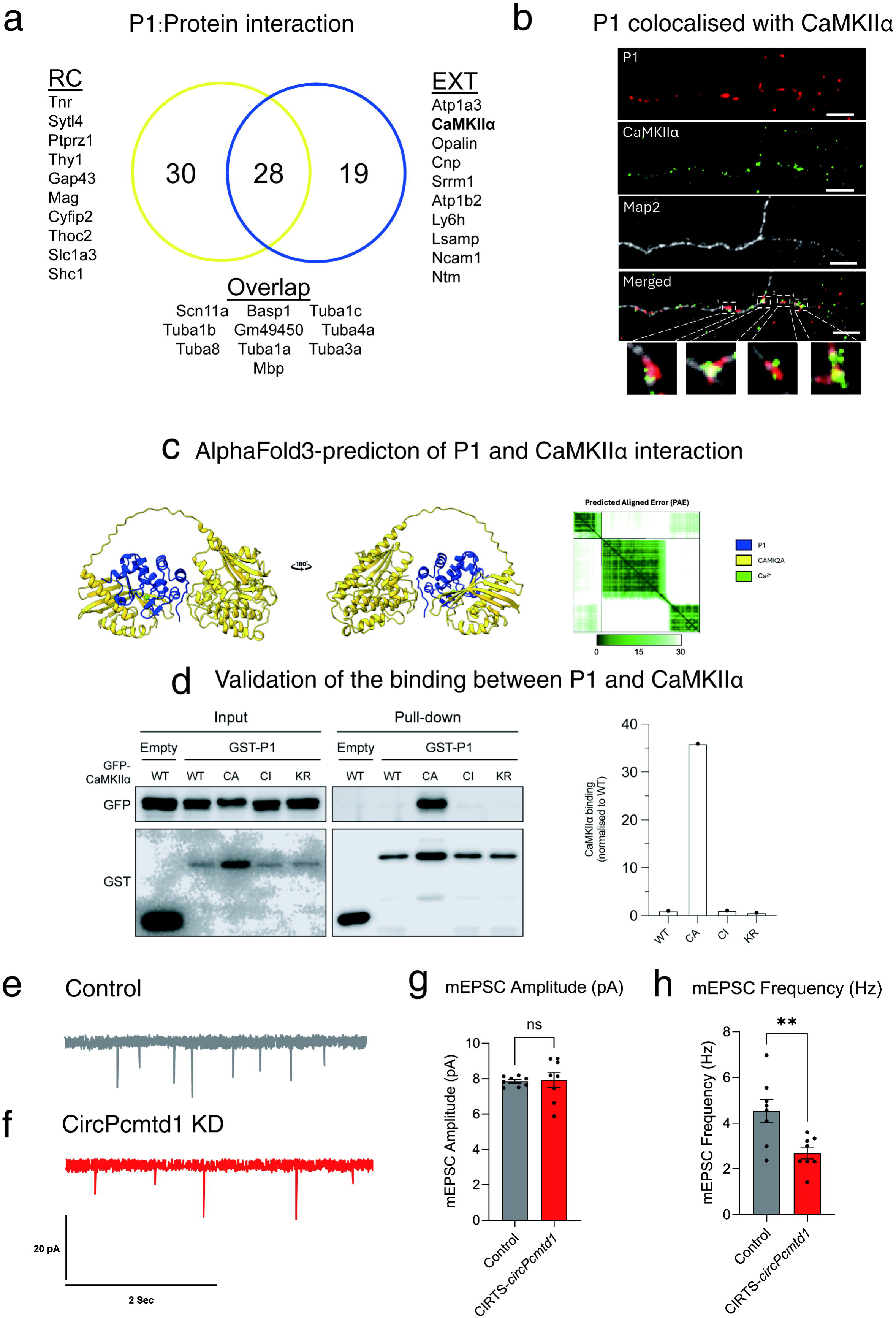
**a.** The Venn diagram shows the number of unique and common proteins that bind to P1 in both the fear conditioned (RC) and extinction-trained (EXT) groups. **b.** Representative confocal image of P1 and CaMKIIa co-localization in dendrites following KCl-induced depolarisation. **c.** AlphaFold3 prediction of the binding between P1 and CaMKIIα. The panel depicts the interaction in the presence of Ca²⁺. P1 is shown in blue, CaMKIIα in yellow, and Ca²⁺ in green. **d.** GST pull-down experiments from HEK293T cells co-transfected with plasmids encoding GST-P1 and GFP-CaMKIIα including WT, Constitutive active (CA), Constitutive inactive (CI), and the K42R, Kinase-dead (KR). Total cell lysates (input) and bound proteins (pull-down) were resolved by SDS-PAGE and analysed by western blotting with specific antibodies against GFP and GST. Bar plot shows the quantification of the levels of P1 binding to GFP-CaMKIIα (WT, CA, CI and KR). Data represent GFP band intensities normalized to WT values. **e.** and **f.** Traces of mEPSCs recorded in primary cortical neurons in the presence of control (Top) and circPcmtd1-Knockdown (Bottom). **g.** and **h.** mEPSC amplitude and frequency of primary cortical neurons in the presence of control and circPcmtd1-Knockdown. n = 8 independent replicates; two-tailed unpaired Student’s t test with Welch’s correction; amplitude, t (20.89) = 0.2017, ns p = 0.8453; frequency, t (20.82) = 3.226, **p = 0.0087. Error bars represent S.E.M.

To provide support for a putative functional interaction between P1 and CaMKIIα, immunofluorescence imaging revealed significant co-localization between the two proteins along Map2-labelled dendrites in activated neurons (Fig. 3b). Given that AlphaFold3 predicted that the relationship between P1 and CaMKIIα, may be calcium sensitive (Fig. 3c), we examined the interaction between P1 and CaMKIIα by co-immunoprecipitation (co-IP) assay and found that there was a high level of binding affinity between P1 and CaMKIIα, specifically when CaMKIIα is in its active state (Fig. 3d). In addition, circPcmdt1 knockdown in primary cortical neurons led to a reduction in miniature excitatory postsynaptic currents (mEPSCs), with no effect on spike amplitude (Fig. 3e-h). Taken together, the data suggest a role for P1 in the regulation of synaptic activity, in part, through a direct interaction with CaMKIIα.

### Targeted knockdown of *circPcmtd1* at the synapse impairs fear extinction memory

Next, to determine whether localised *circPcmtd1* activity is associated with fear-related learning and memory, we used a tool to selectively degrade target circRNAs in the synaptic compartment based on our recent work on RNA localisation and memory^13^ (Fig. 4a). The efficacy of the *circPcmtd1* knockdown construct was first tested *in vitro*, revealing greater then ∼50% reduction in expression, with no significant change in linear Pcmtd1 mRNA levels (Fig. 4b). Next, to confirm the selectivity of targeting *circPcmtd1* at the synapse, we infused the circPcmtd1 knockdown construct into the infralimbic prefrontal cortex (ILPFC) of male mice prior to exposure to a strong (60CS) fear extinction training protocol, which showed a selective reduction in *circPcmtd1* in the synaptic compartment, with little effect on nuclear *circPcmtd1* expression (Fig. 4c) and excellent transfection, *in vivo* (Fig. 4d). Following viral transfection and behavioural training (Fig. 4e), there was no effect of circPcmtd1 knockdown on within-session fear extinction learning or on the ability to express fear when tested 24 hours after training (Fig. 4f and 4g, RC group). In contrast, *circPcmtd1* knockdown led to a significant impairment in fear extinction memory (Fig. 4g). Together, these data suggest that synapse-specific knockdown of *circPcmtd1* in the ILPFC impairs the formation of fear extinction memory without interfering with the original fear memory trace.

**Figure 4.**
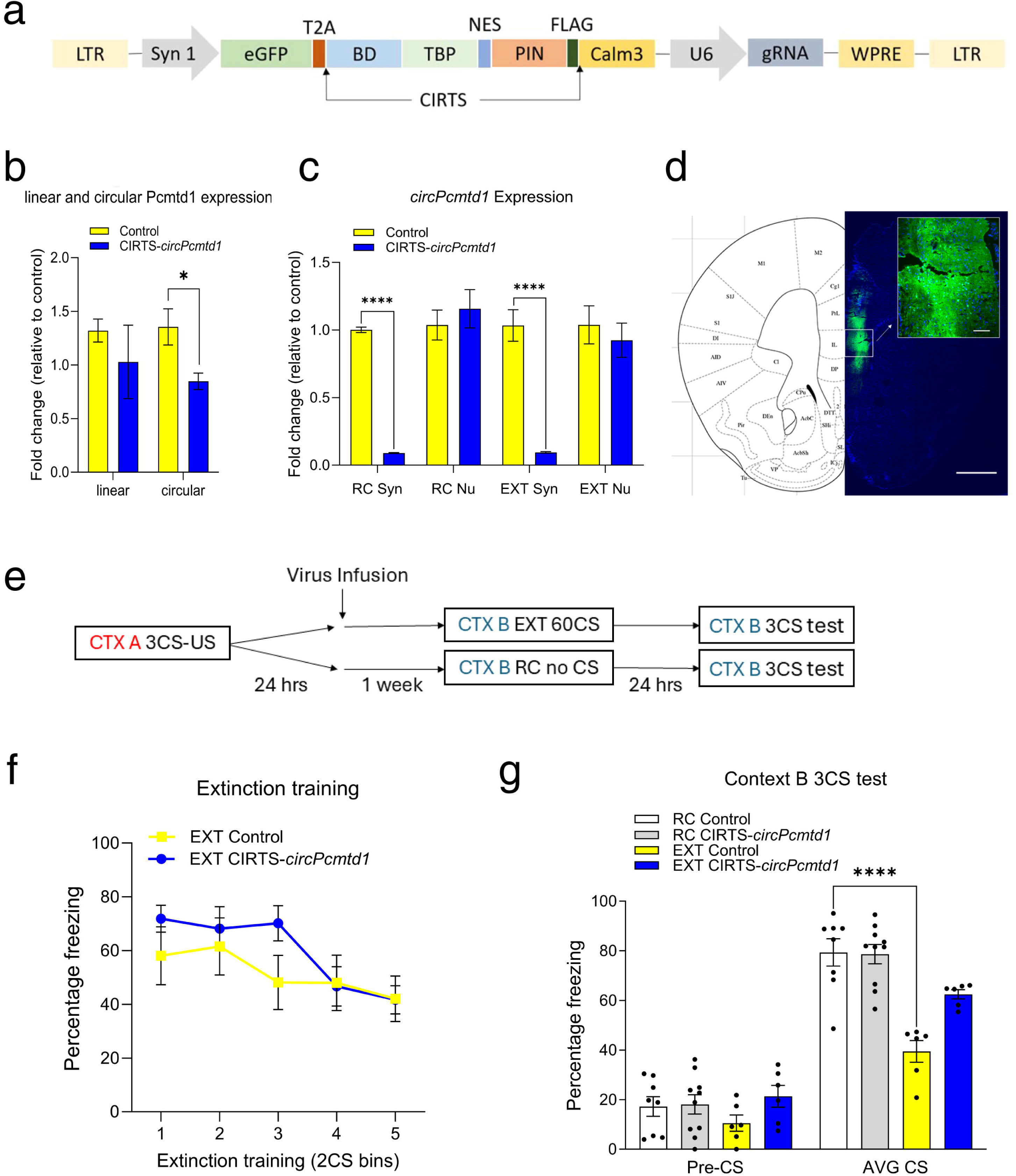
**a.** Schematic of CIRTS-circPcmtd1 construct used for targeted degradation of circPcmtd1. **b.** In vitro validation of CIRTS-circPcmtd1 knockdown in primary cortical neurons showing reduced circPcmtd1 levels without interfering with linearPcmtd1 expression (n ≥ 5 independent biological replicates per group, two-tailed unpaired Student’s t-test, linearPcmtd1 CIRTS control vs. CIRTS-circPcmtd1, t=0.8846, ns p=0.3994; circPcmtd1 CIRTS control vs. CIRTS-circPcmtd1, t=2.553, *p=0.0311). **c.** In vivo validation of CIRTS-circPcmtd1 knockdown in synaptic (syn) and non-synaptic (Nu) lysates isolated from the mPFC of EXT and RC mice. There was significant decrease in circPcmtd1 expression in the synapse for both RC and EXT groups (n=6 biologically independent replicates per group; two-tailed unpaired student’s t-test, RC Syn CIRTS control vs. CIRTS-circPcmtd1, t=44.71, ****p<0.0001; RC Nu CIRTS control vs. CIRTS-circPcmtd1, t=0.6720, ns p=0.5168; EXT Syn CIRTS control vs. CIRTS-circPcmtd1, t=8.038, ****p<0.0001; EXT Nu CIRTS control vs. CIRTS-circPcmtd1, t=0.6000, ns p=0.5618). **d.** Representative image of viral infusion of CIRTS-circPcmtd1 in the ILPFC; scale bars are 1000μm and 100μm, respectively. **e.** A schematic of the behavioural protocol used to test the effect of lentiviral-mediated degradation of circPcmtd1 in the ILPFC on fear-extinction memory. CTX refers to context, CS refers to the conditioned stimulus (tone), and US refers to the unconditioned stimulus (foot shock). **f.** There was no effect of CIRTS-circPcmtd1 knockdown on performance during within-session extinction training (n≥6 biologically independent replicates per group, two-way repeated measures ANOVA, F_1,55_=2.415, p=0.1259). **g.** Relative to mice treated with the control virus that were fear conditioned and exposed to the novel context (RC Control) extinction trained (EXT control) showed a normal reduction in freezing; however, this effect was not present EXT CIRTS-circPcmtd1 mice (n≥6 biologically independent replicates per group, two-way ANOVA, F_3,26_=9.72, p=0.0002; Dunnett’s post-hoc test, RC control vs. EXT control, AVG CS ***p<0.0001, RC control vs. EXT CIRTS-circPcmtd1, AVG CS *p=0.0237, RC control vs. RC CIRTS-circPcmtd1, AVG CS ns p=0.9975). All data are plotted as mean values with error bars representing the SEM.

### P1 overexpression enhances fear extinction memory

To establish a causal role for P1 in fear extinction, we overexpressed P1 using a lentiviral-driven construct (Fig. 5a), which exhibited a robust increase in the mRNA encoding P1 (Fig. 5b) as well as P1 protein levels in, *in vivo* (Fig. 5c) and robust transfection efficiency in the adult mPFC (Fig. 5d). The P1 overexpression construct was then delivered into the ILPFC prior to exposure to a weak (5CS) fear extinction training protocol (Fig. 5e). Limited CS exposure in this manner typically induces no lasting fear extinction memory so it is well suited to examine potential gain of function effects on fear extinction memory. P1 overexpression did not significantly affect extinction learning (Fig. 5f) nor did it have any effect on fear memory per se (Fig. 5g, white vs gray bars). In contrast, P1 overexpression led to a significant reduction in freezing levels in extinction-trained mice compared to all other groups thus indicating a strong gain of function for extinction memory (Fig. 5g). Therefore, these results indicate that increased P1 expression enhances the formation of fear extinction memory.

**Figure 5.**
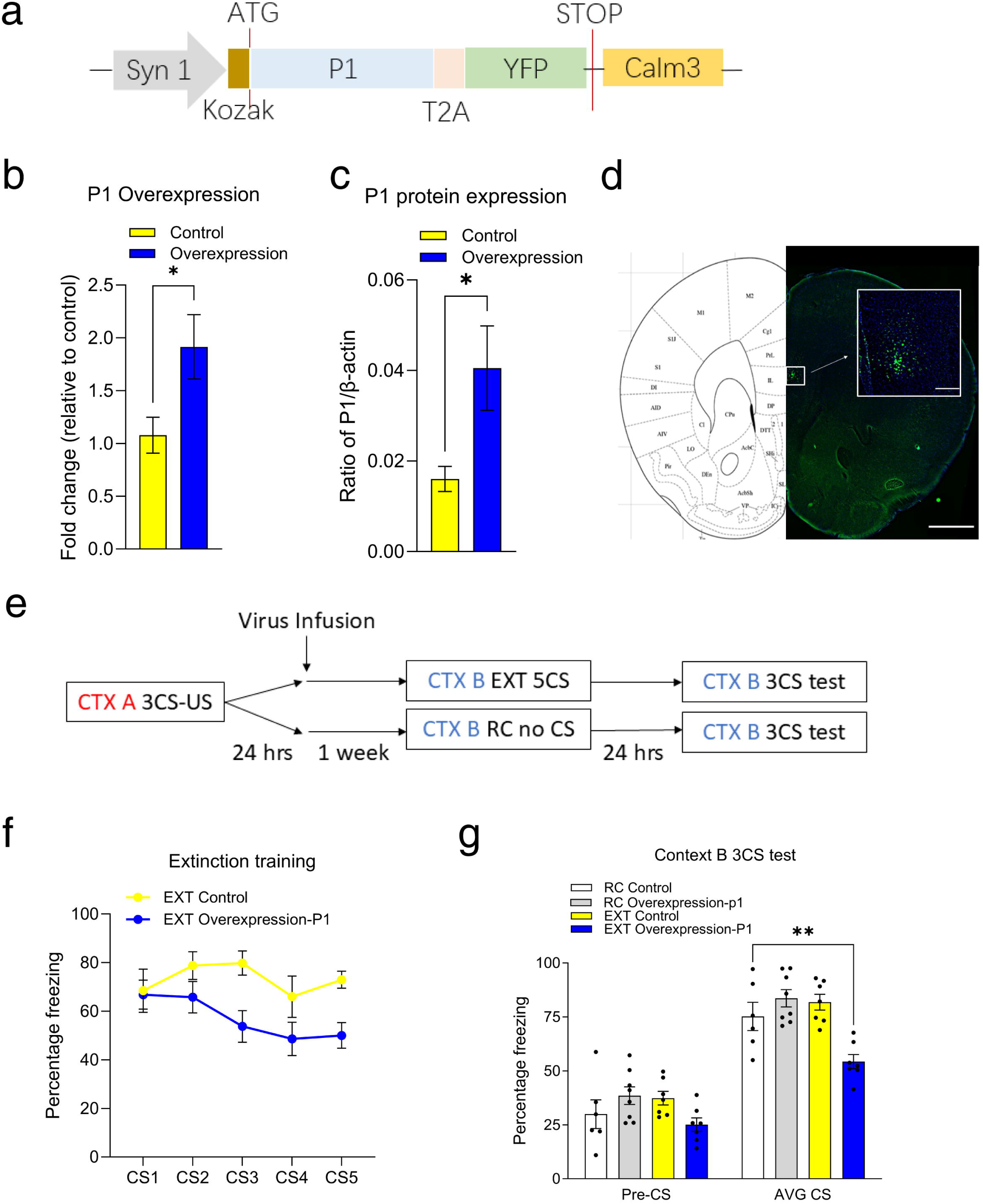
**a.** A schematic of P1-overexpression construct. **b.** *In vivo* validation of overexpression efficiency in the mPFC of EXT overexpression and control mice. There was significant increase in P1 transcript expression (n=6 biologically independent replicates per group; two-tailed unpaired student’s t-test, overexpression control vs. overexpression-P1, t=2.394, *p=0.0377). **c.** Immunoblotting analysis of the expression of P1 in primary cortical neurons following P1 overexpression (n = 4 biological independent replicates per group; two-tailed unpaired student’s t-test, t=2.514, *p=0.0457). **d.** Representative image of P1 overexpression virus in the ILPFC; the scale bars are 1000μm and 100μm, respectively. **e.** A schematic of the behavioural protocol used to test the effect of lentiviral-mediated P1 overexpression on fear-extinction memory. **f.** There was no significant effect of P1-overexpression on within-session EXT training, (n≥6 biologically independent replicates per group, two-way repeated measures ANOVA, F_4,110_=0.6494, p=0.6285). **g.** P1 overexpression had no effect on freezing in mice fear conditioned and exposed to context B only (RC); however, there was a reduction in freezing in EXT trained mice in the presence of P1 overexpression, indicating an enhancement extinction memory that was not seen in mice extinction trained in the presence of control virus. (n≥6 biologically independent animals per group, two-way repeated measures ANOVA, F_3,24_=9.233, p=0.0003; Dunnett’s post-hoc test, RC control vs. EXT control, AVG CS ns p=0.5921, RC control vs. EXT overexpression-P1, AVG CS **p=0.0053, RC control vs. RC overexpression-P1, AVG CS ns p=0.3885). All data are plotted as mean values with error bars representing the SEM.

### P1 exhibits significant enzymatic activity

Having confirmed that the P1 micropeptide accumulates in the synaptic compartment, interacts with synaptic proteins, and is required for memory formation, we next queried whether P1 is a functionally active enzyme. As indicated, P1 is homologous to PIMT, which is a well-known protein methyltransferase (MTase) involved in protein repair via the removal of the post-translational modification isoaspartate. This is potentially important for protein function in the synaptic compartment because the accumulation of isoaspartate leads to protein dysfunction, insolubility, aggregation, and immunogenicity^29^. A recent study found that the host protein Pcmtd1 has lost its MTase activity and instead functions as a chaperone^30^. Thus, we were surprised to find that the P1 micropeptide exhibited very strong MTase activity, comparable to that of PIMT (Fig. 6a-c). Protein methylation influences various biological processes, including gene expression, transcriptional regulation, and signal transduction^31^, and dysregulated MTase activity has been linked to range of neurodegenerative conditions^32^. Therefore, it is plausible that P1 regulates synaptic activity with subsequent effects on memory, in part, through its capacity to target CaMKIIα and other synaptic proteins known to acquire the isoaspartate lesion. Future studies will aim to resolve the precise relationship between P1 MTase activity and the accumulation of isoaspartate on CamKIIα and reveal the temporal role of P1 in CamKIIα stability, function, and repair.

**Figure 6.**
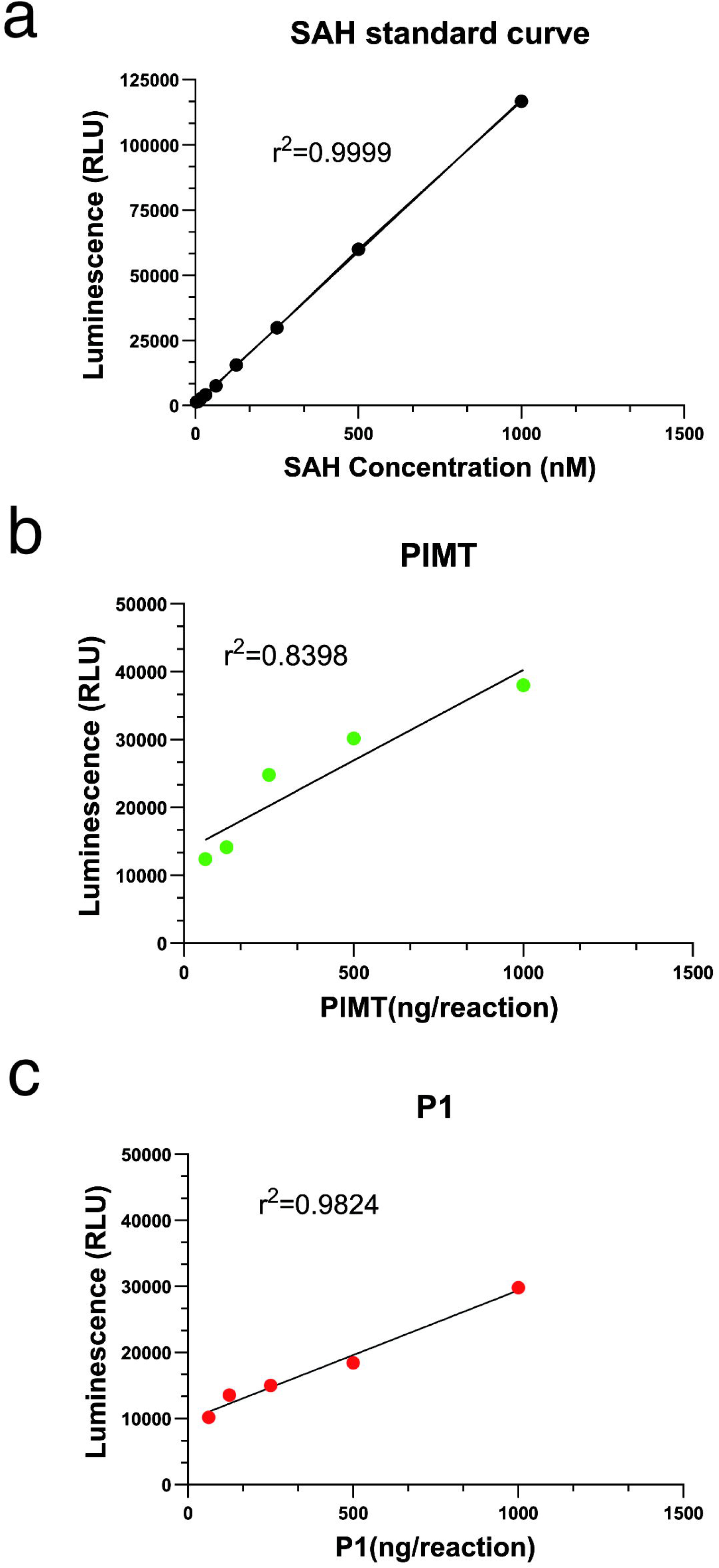
Methyltransferase enzyme activity test. **a.** Representative SAH standard curve r^2^=0.9999. Titration of methyltransferases using the MTase-Glo™ Assay for PIMT and 6xHis-tagged P1 on denatured total mPFC proteins as substrate revealed significant methyltransferase activity in both **b.** PIMT r^2^=0.8398 and **c.** P1 r^2^=0.9824, respectively. Luminescence was measured using a plate-reading luminometer.

## Summary

In this study, we have made three key discoveries surrounding the role of circRNAs in the adult brain: 1) using orthogonal genomics and biochemical approaches, we found widespread experience-dependent circRNA translation in the synaptic compartment, 2) we identify a novel synapse-enriched micropeptide, P1, that directly interacts with synaptic proteins involved in synaptic activity and learning, potentially acting as a protein repair enzyme, and 3) that P1 is required for the formation of fear extinction memory.

The field of micropeptide biology has recently exploded with the realisation that there are thousands of uncharacterised proteins derived from non-canonical sources within the genome, including untranslated regions (UTRs), out of frame internal open reading frames (ORFs), and small proteins derived from non-coding RNAs. For example, there is a surprising number of cytoplasmic long noncoding RNAs that encode novel micropeptides expressed in various tissues, including the brain^33,34^. In addition, it is now evident that circRNAs have the capacity to generate peptides^9^. In line with this, Pamudurti et al used ribosome footprinting and mass spectrometry to reveal a circRNA called *circMbl* that generates a novel peptide in the *Drosophila* brain^35^. Our finding of widespread experience-dependent circRNA translation at the synapse and the discovery of the P1 micropeptide posits a new role for non-canonical proteoforms in the regulation of fear extinction memory.

As indicated, a GO analysis revealed a significant proportion of synapse-enriched circRNAs that potentially encode enzymes capable of protein modification and repair. We verified that P1 is a functional enzyme that is distinct from its full-length host protein Pcmtd1, which has lost its enzymatic capacity^30^. Moreover, we also observed specific interactions between P1 and plasticity-related proteins, which are critical for the regulation of intrinsic plasticity of cortical neurons and that are required for fear extinction memory. Given the important role of CaMKIIα in synaptic plasticity and memory consolidation, the observed interaction between P1 and CaMKIIα agrees with this idea. Moreover, since circPcmtd1 knockdown reduced the frequency of mEPCs, the data support the idea that P1 is functionally involved in the regulation of synaptic activity underlying the formation of fear extinction memory.

We propose that micropeptides generated by the local translation of circRNAs could serve to help maintain optimal functional states of plasticity-related proteins and/or organelles in the synaptic compartment. In the case of P1, the *de novo* generation of a single domain IDR could confer promiscuous protein binding with the specificity of P1 targets determined by the subcellular compartment in which it resides. Akin to the inverse relationship between miniaturization and computational capacity articulated by Moores law, scaling down to make smaller proteins that fit in the synaptic compartment, locally active circRNA-derived micropeptides like P1 would exert their effects with much higher efficiency, faster temporal dynamics, and less energetic demands than those required for the synthesis and trafficking of much larger proteins emerging from the cell body, thereby enabling functional regulation of target proteins at the synapse that would align with the temporal dynamics of learning and memory.

The precise mechanisms governing local circRNA translation remain to be fully characterized. Nonetheless, the evidence suggests IRES-like activity is critically involved in facilitating cap-independent translation of circRNA^9^. Given that the initiation of IRES-mediated translation is favored under conditions of stress, a potential mechanism emerges whereby specific classes of circRNAs may undergo rapid local translation to produce protein modification enzymes with activity comparable to their parental genes, potentially serving as molecular buffers at the synapse in response to stress. In addition, the observation that a significant number of synapse-enriched circRNAs exhibited IRES activity not directly related to translation suggests other regulatory roles for circRNAs in the synapse, including SINEUP and ribozyme activity, which could further impact target RNA translation or other cellular processes.

In conclusion, the discovery of experience-dependent local circRNA translation sheds new light on a previously unknown source of the ‘dark’ proteome in the brain, confirms a critical role for synapse-enriched circRNAs in the regulation of critically adaptive processes, including fear extinction, and suggests a novel mechanism by which neurons have overcome the challenge of space and time for learning and memory.

## Acknowledgments

We gratefully acknowledge grant support from the McCusker Foundation and the National Health and Medical Research Council (GNT2003414), National Natural Science Foundation China (82171517, 822711556, 82471534), and Anusandhan National Research Foundation (CRG/2022/003117). S.U.M. and E.L.Z. were supported by a Westpac Future Scholarship. S.B. and S.S. are supported by a core research grant from the Science and Engineering Research Board from the government of India. We thank Dr. Alun Jones from the Mass Spectrometry Facility in the Institute of Molecular Bioscience at the University of Queensland for assistance with the proteomics experiments, the Queensland Brain Institute Advanced Microscopy Facility and Flow Cytometry Facility for its support.

## Materials and Methods

### Animals

Male C57BL/6J mice (10–12 weeks old) were maintained under standardized housing conditions: four mice per cage, with a 12-hour light/dark cycle, temperature range of 18–24°C, and relative humidity of 30– 70%. Mice were provided *ad libitum* access to standard rodent chow and water. Experimental procedures were conducted during the light phase under red-light illumination, in accordance with protocols approved by the Institutional Animal Care and Animal Ethics Committees of the University of Queensland and Wuhan University.

### Plasmid Construction

The mRuby-ZKSCAN1-split-eGFP reporter plasmid was obtained from Prof. Howard Chang and Dr. Chun-Kan Chen (Stanford University, USA). The CIRTS-*circPcmtd1* vector was prepared by PCR amplification of U6-circPcmtd1 gRNA and U6-scrambled control gRNA, which were then inserted into the XbaI site of the CIRTS vector (Addgene #213172). The Overexpression-P1 vector was constructed by amplifying and inserting *circPcmtd1* ORF-T2A-YFP cassettes into the AgeI and XbaI sites of the pFsy (1.1) GW lentiviral expression vector (Addgene #27232). For the P1 *in vitro* synthesized vector, the coding sequence of P1 was PCR amplified and inserted into the NcoI and XhoI sites of pET His6 TEV LIC cloning vector (Addgene #29653).

### Primary Cortical Neuron Culture

Cortical tissue was isolated from E16–E18 C57BL/6J embryos. Primary cortical neurons were prepared by carefully removing the skull and meninges, followed by tissue dissociation. Cells were resuspended in Neurobasal medium (Gibco) supplemented with 2% B27 supplement (Gibco), 2 mM GlutaMAX (Gibco), and 1% penicillin-streptomycin (Gibco). The cell suspension was homogenized through gentle pipetting and passed through a 40 µm cell strainer (BD Falcon) before plating on poly-L-ornithine (Sigma) coated culture dishes or coverslips (neuVitro). HEK293T and HT22 cell lines were maintained in Dulbecco’s Modified Eagle’s Medium (DMEM) (Gibco) with high glucose, 5% fetal bovine serum (Gibco), and 1% penicillin-streptomycin (Gibco). Media were refreshed every third day.

### Ribo-Seq

Ribosome-bound RNAs were isolated from the mPFC of behaviorally trained Tagger mice. First, naïve 9-week-old male CaMKIIα Cre-Tagger mice (n=14/group) were implanted with a 22-gauge dual guide cannula into the mPFC, after which they were given 1 week to recover. The mice then underwent fear conditioning with 3 pairings of a tone (2 min, 80 Hz) with a foot shock (0.7 mA, 2 s). 24 hours later, they underwent extinction training with 60 non-reinforced tone exposures (EXT) and were sacrificed immediately after fear extinction training. After the mPFC was dissected and synaptosomes isolated by fractionation, RNA immunoprecipitation for ribosome-bound RNA was performed using Rpl22-HA pull down, and 200ng of barcoded RNA was prepared for sequencing (Ribo-Seq). The RNAs were fragmented to 200-500bp and sequencing libraries generated using the SMARTer® Stranded Total RNA-Seq Kit v2 - Pico Input Mammalian (TAKARA #634413) for Illumina. The prepared libraries were then sequenced on the NovaSeq 6000 (2×150bp) (GENEWIZ).

### IRES-Seq

IRES-Seq followed a previously published method^9^. Specifically, an oligo library containing 3,139 oligonucleotides (Twist Bioscience) was prepared based on the identification of 1,819 circRNAs enriched in synapses from our previously published circRNA-seq dataset^7^. These circRNAs were identified as having a synapse-to-nucleus fold-change ≥ 2 and an adjusted p-value < 0.001. We designed 174-nt oligonucleotides using a sliding window approach with a step size of 87 nt to cover the exonic regions flanking the backsplice junction (BSJ). For circRNAs consisting of a single exon, the exon sequence was duplicated and joined end-to-end to position the BSJ centrally, after which the same sliding window strategy was applied to the resulting concatenated sequence. After 5 days of culture, HEK293T cells were sorted into five expression bins based on mRuby and eGFP signal (BD FACSymphony™ S6 Cell Sorter). Approximately 250000 mRuby(+)/eGFP(+) and 20000 mRuby(+) cells were collected for analysis. Total DNA from the respective expression bins was extracted, and three rounds of PCR were conducted to generate sequencing libraries with Illumina-compatible adapters and barcodes. The libraries were first purified using 0.7x volume AMPure XP beads (Beckman) and 2% agarose gel electrophoresis, followed by concentration measurement with Qubit fluorometer (Thermo Fisher Scientific) and quality control using a 2100 Bioanalyzer system (Agilent). Finally, the libraries were sequenced on an Illumina NovaSeq 6000 platform with 150-bp paired-end reads (GENEWIZ), generating ∼15-20 million reads per bin.

### Sequencing analysis

To identify significantly synapse-enriched circRNAs, we reanalyzed previously generated datasets, defining significance as FDR < 0.001 and log₂FC > 1. This analysis yielded 3,285 circRNAs from 70,377 candidates in the RC group and 2,746 circRNAs from 71,139 candidates in the EXT group^7^.

To identify full-length circRNA isoforms, we employed CIRI-Full^36^ and CIRI-vis^37^ with default parameters. For each sample the output of CIRI-Full used custom scripts with mouse genome (mm10) as reference. Resulting isoform-level expression profiles were used for downstream analyses.

For IRES-Seq analysis, adapter and primer sequences were trimmed using Cutadapt (v4.9). The cleaned reads were aligned to the designed IRES oligo library using Bowtie2 (v2.5.4) with default parameters. Reads with a MAPQ score ≥ 3 were retained, to ensure the high-quality alignment, and multi-mappers were excluded. Mapped read counts per oligo were then calculated for each sample.

Circular RNA profiling was also performed using the Ribo-seq data, with CIRIquant (v1.1.2) employed to detect ribosome-bound circular RNAs. This analysis generated count matrices of BSJ reads for 1,819 synapse-enriched circular RNAs across 28 samples, comprising 14 input control samples and 14 IP samples. Raw read counts were imported into R and processed using the edgeR package (v4.4.1) to assess differential expression between IP and input samples. To account for variability in sequencing depth, the total number of sequencing reads per sample was used as the library size. Read depths ranged from 100 million to 229 million reads per sample, with input samples averaging 179 million and IP samples averaging 172 million reads. Low-abundance circular RNAs were filtered out prior to analysis. Only those with more than one BSJ read in at least 14 samples (≥50% of all samples) were retained, reducing the dataset from 1,819 to 1,450 circular RNAs for downstream analysis. Differential expression analysis was performed using the edgeR workflow. Common and tagwise dispersions were estimated with the estimateDisp() function, using a design matrix that accounted for sample type (Input vs IP). Quasi-likelihood F-tests were then conducted using the glmQLFit() and glmQLFTest() functions to identify circular RNAs enriched in IP samples relative to input. P-values were adjusted for multiple testing using the Benjamini–Hochberg false discovery rate (FDR) method.

For the analysis of B2 SINE sequences, we first identified circRNAs that were enriched in Ribo-seq data and had corresponding full-length predictions (FDR < 0.05, Log₂FC > 0.585). Among 842 circRNAs enriched in Ribo-seq, 300 isoforms were found to have full-length predictions. We then extracted the genomic coordinates of these circRNAs and converted them into BED format using a custom R script. The corresponding sequences were retrieved from Mouse mm10 genome using the Galaxy Bedtools getfasta tool (Galaxy Version 2.31.1+galaxy). Finally, we applied a custom R script to detect the presence of *GGUGGU* and *GUGGC* motifs within the extracted sequences.

For the ORF prediction on the circRNAs, their full sequences were reconstructed from genomic coordinates by concatenating the backsplice junction ends into linear forms. ORFs were predicted using TransDecoder (v5.5.0), and the predicted peptides were cross-referenced against mass spectrometry (MS) data. ORFs supported by MS evidence were considered high confidence.

### Synaptosome Preparation

Synaptosome preparation followed the Syn-PER™ Synaptic Protein Extraction Reagent (Thermo Scientific #87793) protocol. The mPFC of one to four mice was homogenized in a buffer containing 1ml Syn-PER Reagent, 5 mM DTT, 0.1 mM RNaseOUT (Invitrogen) or Protease Inhibitor Cocktail 100x (Thermo Scientific) using a Teflon-glass tissue grinder. The homogenate underwent centrifugation at 1,000 × g for 10 minutes at 4°C, with the nucleus-enriched pellet retained for subsequent analysis. The supernatant was further centrifuged at 15,000 × g for 20 minutes at 4°C, yielding a synaptosome pellet suitable for RNA or protein extraction.

### RNA Extraction

Cultured cells and tissues were homogenized using a dounce tissue grinder. Homogenate lysis or synaptosomes were then digested using NucleoZOL (Macherey-Nagel) supplemented with 5 mM DTT (Thermo Scientific) and 0.1 mM RNaseOUT (Invitrogen). Samples were centrifuged for 15 minutes at 12,000 × g. The RNA-containing supernatant was precipitated with 100% isopropanol and purified using 70% Ethanol. RNA concentration was measured using a nanophotometer (IMPLEN) or Qubit fluorometer (Thermo Fisher Scientific).

### RT-qPCR

cDNA synthesis was performed using 1 µg of RNA with either the QuantiTect Reverse Transcription Kit (Qiagen) or SensiFAST cDNA Synthesis Kit (Cat. No. BIO-65053, Bioline) according to the manufacturer’s instructions. Quantitative PCR was performed on a RotorGeneQ real-time PCR cycler (Qiagen) with SensiFAST SYBR master mix (Cat. No. BIO-98005, Bioline) using primers for target genes. Transcript levels were normalized to β-Actin using the ΔΔCT method, with each PCR reaction run in duplicate and repeated at least twice per sample.

### Protein Extraction

Cultured cells and homogenized tissue were lysed in cell lysis buffer II (Invitrogen) containing 100x Halt Protease Inhibitor Cocktail (Thermo Fisher Scientific) and DTT (Cat. No. R0861, Thermo Fisher Scientific). Specifically, for the P1-CaMKIIα or p-CaMKIIα co-IP experiment, the cells were Lysed in cell lysis buffer II (Invitrogen) supplemented with Complete protease inhibitor cocktail (Sigma-Aldrich) and PhosSTOP (Sigma-Aldrich). Samples were incubated on ice for 30 minutes, then centrifuged at 13,000rpm for 10 minutes at 4°C. Supernatant protein concentration was measured using the Bradford assay (Sigma) on a PHERAstar Microplate Reader (BMG LABTECH). Protein samples were diluted in Laemmli buffer with 5% 2-mercaptoethanol (Sigma-Aldrich) and denatured at 95°C for 5 minutes.

### Immunoprecipitation (IP)

Antibody-bead conjugation was performed by incubating 5–10 µg of antibody with 10 µl of Protein G (Invitrogen) or Protein A (Invitrogen) slurry in 200 µl of PBST for 1 hour. After centrifugation at 2,400 × g for 5 minutes, the supernatant was discarded. The antibody-bound beads were then treated with 200 µl of blocking buffer for 30 minutes to minimize non-specific interactions. Following washing with PBST, 100– 200 µl of prepared lysate was added to the beads and incubated at 4°C for 1–2 hours with continuous rotation. Post-protein capture, beads were pelleted by centrifugation, and the supernatant was retained for downstream analysis. Immunoprecipitated complexes were washed twice with PBST containing 1% NaCl, centrifuging at 2,400 × g for 5 minutes after each wash. For SDS-PAGE analysis, the final bead pellet was resuspended in 4x Laemmli SDS sample buffer (Alfa Aesar) and 1 µl 2-mercapethonal (Sigma). After boiling at 95°C for 5 minutes, the sample was directly loaded onto the polyacrylamide gel.

### Binding assay

HEK293T cells were co-transfected with P1-GST together with one of GFP-CaMKIIα (WT), GFP-CaMKIIα (Constitutive active), GFP-CaMKIIα (Constitutive inactive), or GFP-CaMKIIα (K42R, Kinase-dead) in combination by the calcium-phosphate precipitation method. After 48 h, cells were harvested and lysed in an ice-cold buffer containing 1% Triton X-100, 1 mM EDTA, 1 mM EGTA, 50 mM NaF, and 5 mM sodium pyrophosphate in PBS, supplemented with an EDTA-free protease inhibitor cocktail. The lysates were centrifuged at 20,627 × g for 20 min at 4 °C, and the supernatant was incubated overnight at 4 °C with glutathione agarose beads (Thermo Scientific). Beads were washed three times with cold lysis buffer, and the bound proteins were released with 2× SDS loading buffer at 50 °C for 30 min followed by western blotting. Blots were incubated with mouse anti-GFP (Cat no. ab1218; mouse monoclonal; Abcam), mouse anti-GST (Cat no. 66001-2-Ig; mouse monoclonal; Proteintech) antibody, respectively, and then were analyzed using the enhanced chemiluminescence method. Images were acquired on the Odyssey Fc imaging system (LI-COR) and band intensities were quantified using Image Studio Lite software (LI-COR).

### Western Blot

Total proteins or IP were resolved by SDS-PAGE, transferred onto PVDF membranes (BioRad), and blocked with Odyssey Blocking Buffer (Li-COR) for 1 hour at room temperature. The membranes were then washed 3 times with PBST. Then the membranes were probed with custom-made anti-P1 (Cat. no. AB014750; rabbit polyclonal; 1:100; BIOMATIK), anti-β-Actin antibody (Cat. no. 8H10D10; mouse monoclonal; 1:1000; Cell Signaling Technology) overnight at 4°C on a shaker. Antibodies were diluted with PBST. The membranes were washed 3 times with PBST for 10 minutes, followed by incubation with secondary antibodies (IRDye 800CW Donkey anti-mouse IgG (H+L), 1:20000 and IRDye 800CW Donkey anti-rabbit IgG (H+L), 1:20000; Li-COR) for 1 hour at room temperature, washed 3 times with PBST, and imaged using an Odyssey Fc system (Li-COR).

### Protein Purification

The P1-pET His6 TEV LIC vector was transformed into *E. coli* strain C3013 (NEB #C 3013I T7 Express lysY/Iq Competent Cells, High Efficiency). Transformed cells were incubated overnight at 37°C. For the initial culture, five single colonies were picked and inoculated into LB medium supplemented with kanamycin, followed by overnight shaking incubation at 37°C. For the secondary culture, each sample was prepared by a 1:20 dilution of the overnight culture and incubated with shaking at 37°C until the optical density at 600 nm (OD₆₀₀) reached approximately 0.6–0.8. Protein expression was induced by adding IPTG (Goldbio) to a final concentration of 1 mM, followed by shaking incubation at 37°C for 3 hours. Cells were harvested by centrifugation, and the pellets were resuspended in lysis buffer (50mM NaH_2_PO_4_, 300Mm NaCl, 10mM imidazole, 1% Tween20, 50% Glycerol, pH 8.0; 2–5 mL per gram of pellet). Lysozyme (1 mg/ml) (Merck) was added, and the suspension was incubated on ice for 30 minutes. Samples were then transferred to 50 mL Falcon tubes and sonicated on ice in 1-minute intervals to prevent overheating, until no detectable odor was present. RNase A (10 µg/ml) (New England Biolabs) and DNase I (5 µg/ml) (Sigma-Aldrich) were then added, followed by incubation on ice for 15 minutes. The lysate was clarified by centrifugation at 10,000 × g for 30 minutes at 4°C, and the supernatant was collected. For affinity purification, 50 µl of HisPure Ni-NTA Resin (Thermo Scientific) was added to each tube, and samples were incubated with gentle shaking at 4°C for 2 hours. The mixture was centrifuged at 4,000 × g for 2–5 minutes at 4°C, and the supernatant (flow-through) was carefully removed, ensuring minimal loss of resin. The resin was resuspended in 500 µl wash buffer (50mM NaH_2_PO_4_, 300Mm NaCl, 40mM imidazole, pH 8.0) and transferred to a 2 mL microcentrifuge tube. After centrifugation at 4,000 × g for 5 minutes at 4°C, half of the supernatant was removed. Subsequently, 1 ml of wash buffer was added, and the suspension was shaken at 4°C for 5–10 minutes, followed by centrifugation at 4,000 × g, 2–5 minutes, 4°C. This wash step was repeated five times. For elution, after the final wash, as much wash buffer as possible was removed. Then, 100–200 µl of elution buffer (50mM NaH_2_PO_4_, 300Mm NaCl, 500mM imidazole, pH 8.0) was added, and the resin was shaken at 4°C for 5–10 minutes before centrifugation at 4,000 × g for 5 minutes, 4°C. At each step, an appropriate volume was collected for SDS-PAGE analysis to assess purity and accuracy. The supernatant was collected, and this step was repeated three times. Desalting and protein recovery were performed using Zeba™ Spin Desalting Columns (Thermo Scientific #89882) according to the manufacturers instructions.

### Methyltransferase activity

The methyltransferase activities of P1 and PIMT were tested using MTase-Glo™ Methyltransferase Assay kit (Promega). For each reaction, de-natured mPFC lysate (95°C boiling) was used as substrate. All samples were processed following a serial twofold dilution with reaction buffer in a Falcon 96-well plate in the presence of 1 µM S-adenosyl-methionine; with the last well as no-enzyme control. Each reaction was incubated at 30°C for 1 hour, and then 5 µl 5x MTase-Glo reagent and 25 µl detection buffer were added sequentially, shaking for 1-2 minutes and incubating at room temperature for 30 minutes, respectively. Both protocols were processed as described in the MTase-Glo assay protocol section 4.A. Finally, luminescence was measured using the CLARIOstar PLUS enzyme microplate reader (BMG LABTECH).

### Lentiviral Production

Plasmid DNA was co-transfected with pMD2.G (Addgene #12259), pRSV-Rev (Addgene #12253), and pMDLg/pRRE (Addgene #12251) into HEK293T cells at 80% confluence using Lipofectamine 3000 (Thermo Fisher Scientific). Sodium butyrate (Sigma) was added 4 hours post-transfection to enhance viral production. After 48 hours of incubation at 37°C and 5% CO_2_, the virus was harvested by ultracentrifugation, and titer was measured using a Lenti-X qRT-PCR titration kit (Clontech).

### Lentiviral Infusion

Lentiviral infusion followed established protocols. Double cannulae (PlasticsOne) were implanted into the ILPFC 7 days prior to viral infusion. Injection coordinates were set at +1.85 mm (AP) and -2.5 mm (DV) from Bregma. Lentivirus (4 µl total) was injected at 0.2 µl/min via two injections, 48 hours apart. Mice were fear-conditioned 24 hours before the infusions and extinction-trained one week post-infusion.

### Behavioral Training and Analysis

Behavioral testing followed our previously published protocol^7,11–13^, using two contexts (A and B). Behavioural recordings in the two conditioning chambers (Coulbourn Instruments) were captured using ceiling-mounted digital cameras and automatically scored using the FreezeFrame software. Fear conditioning in context A which scented with a lemon-alcohol solution (5% lemon and 10% ethanol) involved 120 seconds of exposure followed by three pairings of an 80 dB white noise conditioned stimulus (CS) and a 1s, 0.7 mA foot shock (US). 120 s intertrial interval (ITI) separated each CS. Mice were matched into treatment groups based on freezing scores during the third CS pairing. Extinction training in context B was performed with a spray of vinegar. Mice were first allowed to acclimate for 120 s and followed the extinction protocol consisting of non-reinforced 120 s CS presentations (60CS for knockdown and 5CS for overexpression experiments), each separated by a 5 second ITI. In behavioural control experiments (retention control, RC), animals were exposed to the same context but did not receive any CS presentations. Memory retrieval was assessed 24 hours later by re-exposing the animals to Context B, during which three 120 s CS presentations were delivered, each separated by a 120 s ITI. Memory performance was quantified as the percentage of time spent freezing during the CS presentations.

### Immunofluorescence

Primary cortical neurons were fixed with 10% neutral buffered formalin (Sigma-Aldrich) for 30 minutes, washed in PBS, and incubated with 4% the serum of primary antibody host species in PBST for 1 hour at room temperature. Cells were then incubated with primary antibodies (custom-made anti-P1 (Cat. no. AB014750; rabbit polyclonal; 1:100; BIOMATIK), anti-pCaMKIIα (Cat. no. MA1-047; mouse monoclonal; 1:2000; Thermo Fisher Scientific), anti-CaMKIIα (Cat. no. MA1-048; mouse monoclonal; 1:2000; Thermo Fisher Scientific), anti-MAP2 (Cat. no. ab5392; Chicken polyclonal; 1:2000; Abcam), and anti-PCMT (Cat. no. 60172-1-1g; mouse monoclonal; 1:1000; Proteintech)) overnight at 4°C, washed 3 times with PBST, and incubated with secondary antibodies for 1 hour at room temperature. Cells were stained with 4′,6-diamidino-2-phenylindole (DAPI) (Thermo Fisher Scientific) for 10 min at room temperature, mounted on microscope slides with ProLong Gold Antifade Mountant (Thermo Fisher Scientific), and imaged using an Axio Imager Z1 microscope (Carl Zeiss).

For tissue sections, animals were transcranial perfused with 4% paraformaldehyde, and brains were subsequently immersed in 30% sucrose for cryoprotection prior to sectioning. Target brain sections were cut at a thickness of 14 µm using a Zeiss Microm HM560 cryostat and mounted onto SuperFrost Plus slides (Thermo Fisher Scientific). The slides were then baked at 60 °C for 30 min, washed and incubated with PBST for 10 min at room temperature. Slides were then incubated for 1 hour in blocking buffer and incubated with primary antibody (anti-GFP, abcam, ab6556, 1:2000) at 4 °C overnight. Slices were washed 3 times with PBST followed by adding secondary antibodies. Slices were then incubated at room temperature for 1 hour, washed 3 times with PBST, and incubated with DAPI (Thermo Fisher Scientific) for 10 min at room temperature. Coverslips were applied with Vectamount Permanent Mounting Medium (ACD). Imaging was conducted using a spinning-disk confocal microscope (Marianas system, 3i, Inc.) comprising an Axio Observer Z1 inverted microscope (Carl Zeiss), a CSU-W1 spinning-disk head (Yokogawa Corporation of America), and an ORCA-Flash4.0 v2 sCMOS camera (Hamamatsu Photonics), equipped with 20×/0.8 NA Plan-Apochromat and 40×/1.2 NA C-Apochromat objectives. Images were acquired using SlideBook software, version 6.0 (3i, Inc.).

### Image Acquisition and Analysis

Image acquisition was performed using ZEN 2012 software (Carl Zeiss), with analysis conducted using ImageJ version 1.52p and figure construction utilizing the FigureJ plugin version 1.36.

### HPLC/MS-MS/MS, mass spectrometry and protein Identification

The magnetic affinity beads were digested with 40 μl of trypsin solution (40 ng/μl in 50 mM ammonium bicarbonate buffer, pH 8, Promega) at 37°C in incubator for overnight. Following digestion, the trypsin solution from each sample was collected to a new clean Eppendorf tube separately. The beads were then treated with 200 μl of 5% formic acid/acetonitrile mixture (3:1 v/v) for 30 minutes at room temperature with shaking. Both the trypsin solution and extraction supernatant were combined and dried using vacuum centrifugation. Prior to analysis, samples were reconstituted in 15 μl of 1.0% TFA, vortexed, and briefly sonicated. Peptide analysis employed a microflow HPLC system containing Eksigent, Ekspert nano LC400 uHPLC (SCIEX), Triple TOF 6600 mass spectrometers (SCIEX), and a micro-Duo IonsSpary, ion source. A 5 μl sample aliquot was initially injected on a 5 mm × 300 μm, C18 3 μm trap column (SGE) at 10 μl/min for 6 minutes. The samples were subsequently washed to the 300 μm × 150 mm Zorbax 300SB-C18 3.5 μm column (Agilent) at 45°C with a flow rate of 3 μl/min. Separation utilized a multi-stage gradient with 3 μl/min: initially 2-25% solvent B over 60 minutes, followed by 25-35% solvent B for 13 minutes, and finally 35-80% solvent B in 2 minutes. The column was re-equilibrated at 2% solvent B between injections. Mobile phase composition consisted of 0.1% formic acid in water (solvent A) and 0.1% formic acid in acetonitrile (solvent B). The equipment parameters were set as: 5500 V of ionspray voltage, 80 V of declustering potential, 25 of curtain gas flow, 15 of nebulizer gas 1, 30 of heater gas 2, and 150°C of interface heater. Data acquisition utilized Information Dependent Acquisition mode with 250 ms full-scan TOF-MS acquisition (m/z 350-2000) followed by up to 30, 50 ms product ion scans (ms/ms, m/z 100-1500), using rolling collision energy. Selection criteria for fragmentation included signal threshold exceeding 150 counts and charge states between +2 and +5. Data acquisition and initial processing were conducted with Analyst software (version 1.7) (SCIEX), while protein identification was performed using Protein Pilot (version 5.0) (SCIEX) for database searching.

### Proteomics Data and GO Analysis

Protein Network analysis was conducted using STRING protein query database version 11.5 for *Mus musculus* (https://string-db.org/). Confidence score cut-off was set at 0.7, with p-values corrected using the Benjamini-Hochberg procedure. GO analysis for protein class was first performed with PANTHER using the parent genes of circRNAs with detected unique peptide (https://pantherdb.org/). The enrichment p-values for each protein class are calculated using the hypergeometric distribution, and the Benjamini-Hochberg method is applied to adjust the p-values for false discovery rate control. Pfam analysis was performed using default parameters from EMBL-EBI with the peptide sequences of *circPcmtd1*(https://www.ebi.ac.uk/interpro/). The biological functions of conserved domains were determined using the annotations from NCBI Conserved Domains database.

### Statistical Analysis

Statistical analyses were conducted using GraphPad Prism version 10. Welch’s t-test was used for two-group comparisons, and one-way ANOVA with Dunnett’s test for group comparisons relative to control. Behavioral data were analyzed using two-way ANOVA with post hoc comparisons performed using Dunnett’s test.

---

**Supplementary Figure 1.**
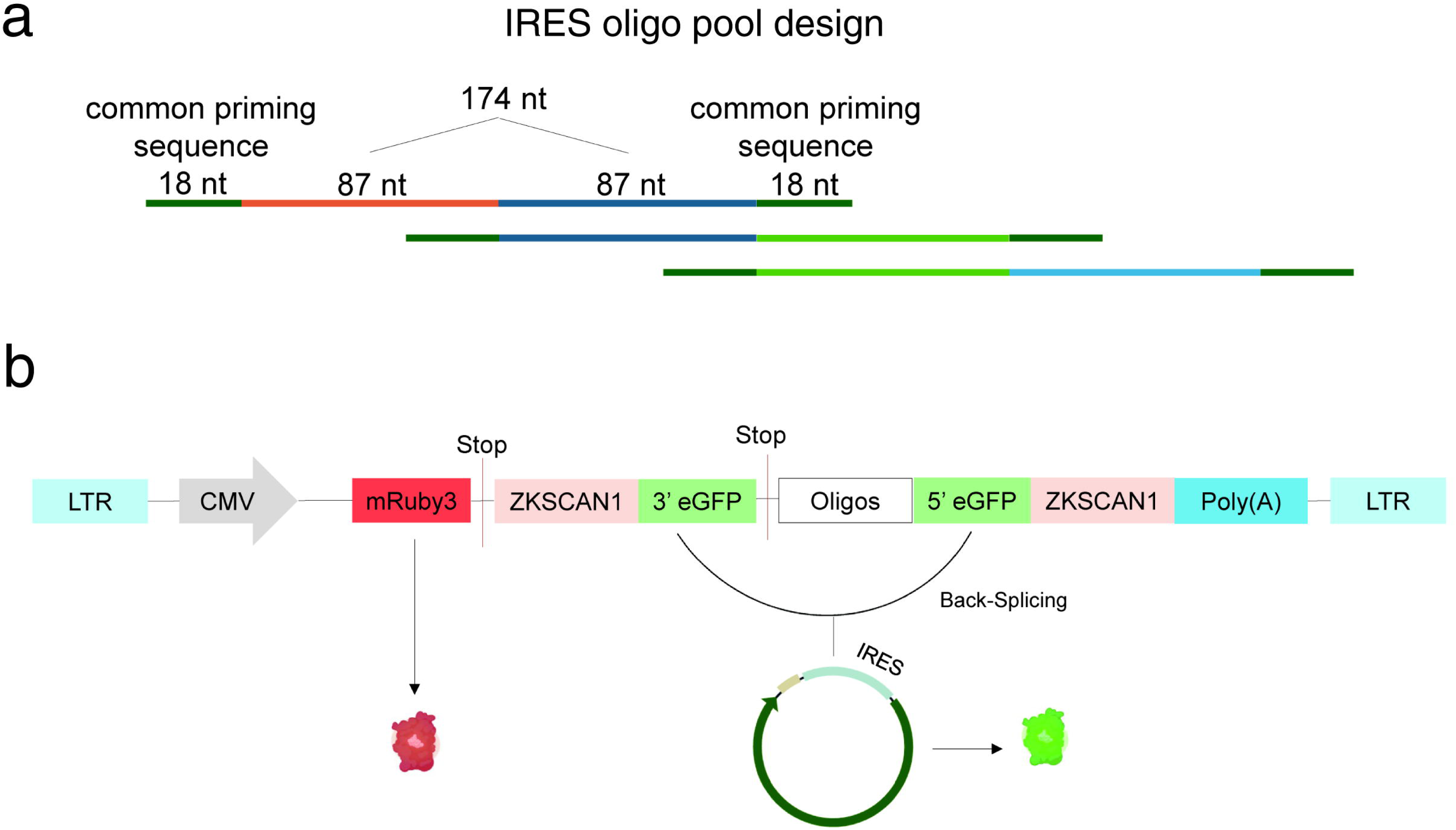
**a.** Diagram showing oligo design using a sliding window approach. **b.** Schematic overview of the high-throughput split-eGFP circRNA reporter screening assay used to identify circRNA IRES.

**Supplementary Figure 2.**
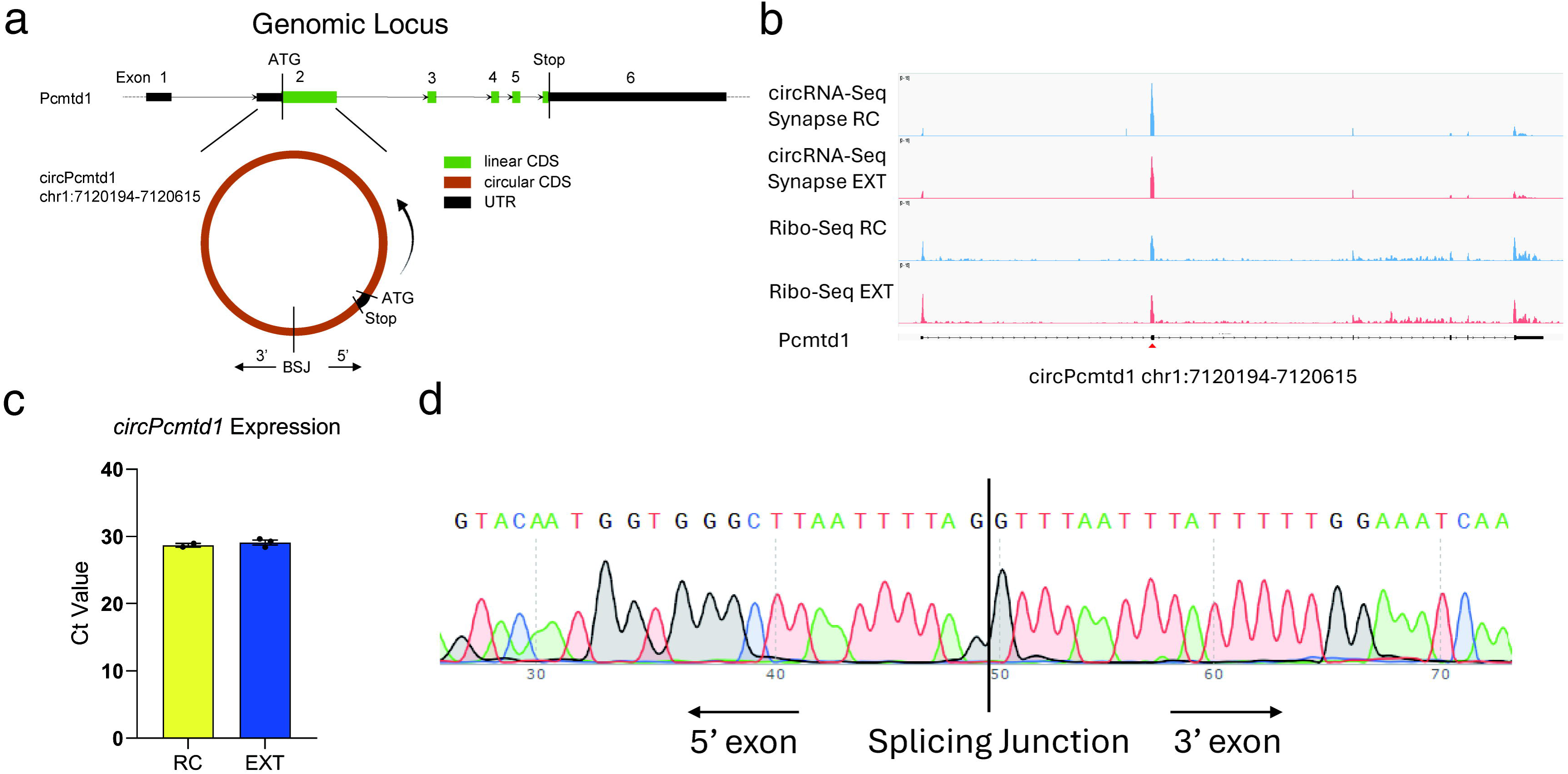
**a.** A schematic of the Pcmtd1 transcript with a total of 6 exons. The CDS region starts from exon 2 to exon 6. *circPcmtd1* is derived from exon 2. UTR: untranslated regions; CDS: coding sequence. **b.** Genomic tracks of the Pcmtd1 gene locus. Bars show BSJ and Ribo-seq read distribution at the synapse from the mPFC of RC (blue) and EXT (red) trained mice. **c.** circPcmtd1 expression in mPFC samples from fear extinction-trained mice. RT-qPCR was performed on RC and EXT mPFC samples. Bar plot based on at least 4 biological replicates. Statistical significance was determined using two-tailed unpaired Student’s t-test. ns, p>0.05. **d.** Sanger sequencing for RT-qPCR products across the BSJ of circPcmtd1.

